# Functional connectivity models for decoding of spatial representations from hippocampal CA1 recordings

**DOI:** 10.1101/073759

**Authors:** Lorenzo Posani, Simona Cocco, Karel Jezek, Rémi Monasson

## Abstract

Hippocampus stores spatial representations, or maps, which are recalled each time a subject is placed in the corresponding environment. Across different environments of similar geometry, these representations show strong orthogonality in CA3 of hippocampus, whereas in the CA1 subfield a considerable overlap between the maps can be seen. The lower orthogonality decreases reliability of various decoders developed in an attempt to identify which of the stored maps is active at the moment. Especially, the problem with decoding emerges with a need to analyze data at high temporal resolution. Here, we introduce a functional-connectivity-based decoder, which accounts for the pairwise correlations between the spiking activities of neurons in each map and does not require any positional information, *i.e.* any knowledge about place fields. We first show, on recordings of hippocampal activity in constant environmental conditions, that our decoder outperforms existing decoding methods in CA1. Our decoder is then applied to data from teleportation experiments, in which an instantaneous switch between the environment identity triggers a recall of the corresponding spatial representation. We test the sensitivity of our approach on the transition dynamics between the respective memory states (maps). We find that the rate of spontaneous state shifts (flickering) after a teleportation event is increased not only within the first few seconds as already reported, but this instability is sustained across much longer (*>* 1 min.) periods.

## 1 Introduction

Over the recent decades, multi-cell recording techniques have provided insights into the nature of brain representations and their internal dynamics. While many works have focused on the input-output transfer functions in primary sensory systems (visual, olfactory, etc.), understanding functions corresponding to complex representations in higher cortical circuits is very hard as they are often based on mixed selectivities (Rigotti et al., 2013). In relatively rare cases, such as in the entorhino-hippocampal system, a highly processed neural activity can be reliably correlated with behavior. The so-called ‘place cells’ in the CA1 and CA3 of hippocampus exhibit sharp spatially tuned and environment specific activity (O’Keefe and Dostrovsky, 1971), see Fig. 1. Collective activity of the place–cell population coding for the environment defines its neural representation, or *map*. Simultaneous recording of multiple place-cell activity thus allows one to identify a general memory state of the network (specific map), as well as to decode the accurate position of the rodent in the corresponding environment (Zhang et al., 1998).

**Figure 1:**
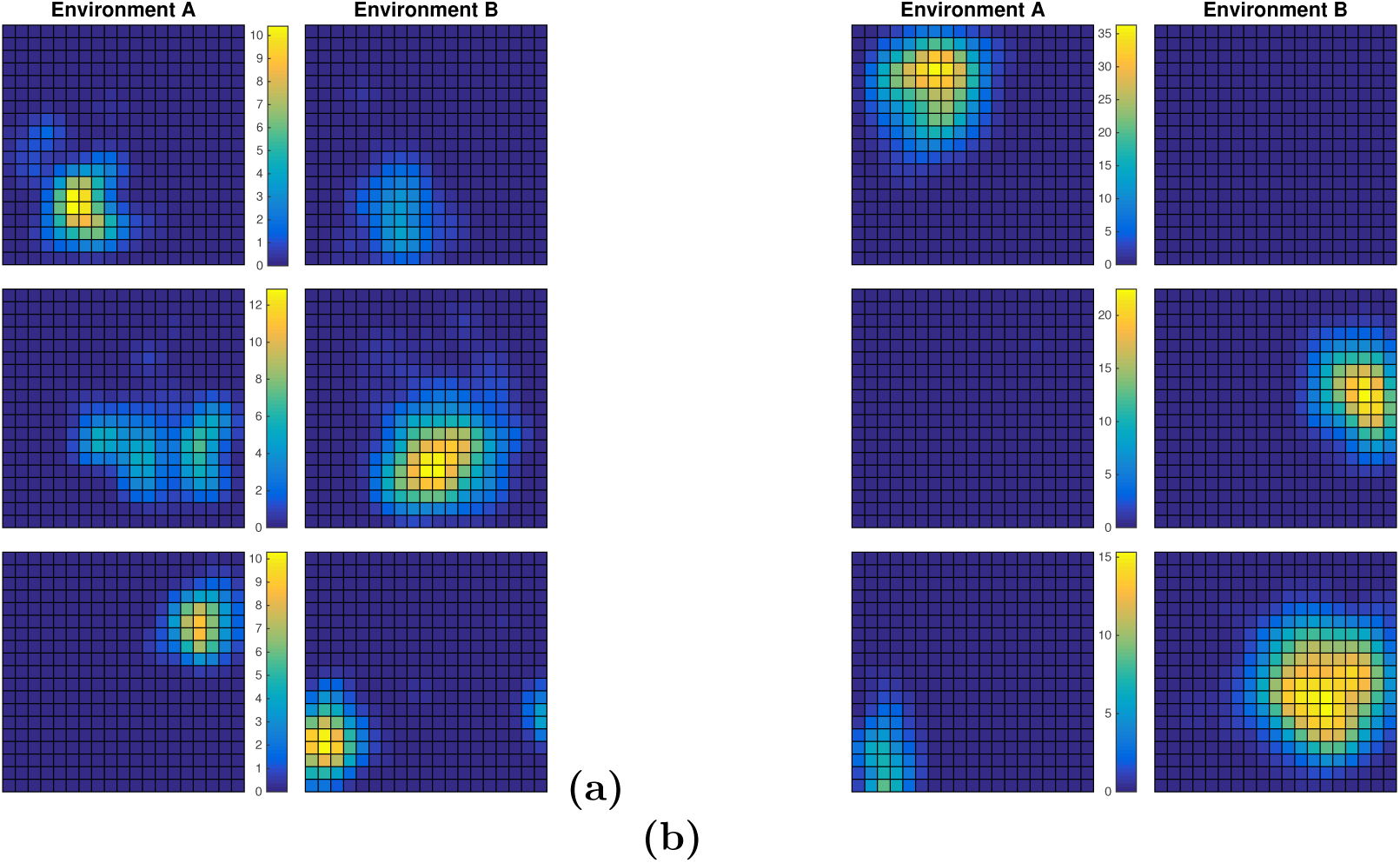
Hippocampal representations are less orthogonal in CA1 (a) than in CA3 (b). Each panel shows six firing fields from CA1 (a) and CA3 (b) corresponding to three place cells (rows) in the recorded neuronal population, computed from 10 min recordings of the activity during free exploration of environments A and B (same 60 *×* 60 cm square shapes; spatial bins : 3*×*3 cm). Whereas CA3 coding is highly sparse and representations are largely orthogonal, CA1 population shows higher amount of cells active in corresponding locations across the two rooms, with peak rates (color scale) changing from one environment to the other. The non-orthogonality of environment representations in CA1 makes identification of the represented map from neural activity difficult compared to the situation in CA3. CA3 data were taken from Jezek et al. (2011). Colorbars show average firing rate in Hz.

Recently, Jezek et al. (2011) have studied the dynamics of transient change between the spatial maps encoding two different environments in CA3 at high temporal resolution (ca 120 ms time windows). The two environments differed by light cues that could be switched instantaneously (‘teleportation procedure’), while the animal hippocampal neural activity was recorded to monitor the course of activation of the proper spatial map. An unstable state generally emerged for some seconds after the light switch, as both maps started to flicker back and forth. This phenomenon, called flickering, was identified through measure of the similarity between the place-cell population activity and its averaged patterns across both environments, recorded earlier in respective reference sessions. Typically, a given 120 ms time window activity of the test data strongly correlated with the average reference activity in one map, and had essentially no correlation with the reference activity in the other map.

Success of such comparison-based decoding methods reflects the strong orthogonality of spatial maps in CA3: across two environments, activity of place cells broadly differ in their mean frequencies and receptive field locations, see Fig. 1(b). Hence, simple map decoders, essentially assuming that cells fire independently of each other, are sufficient to reliably identify the representation expressed by the animal. In contrast, remapping between environments (especially of similar geometry) is less orthogonal in CA1 as it shows higher number of cells firing at corresponding places across rooms, see Fig. 1(a). The population activity vector often correlates well with both concurrent reference templates, which hinders the use of comparison-based methods for map decoding.

Here we address the challenging goal of map decoding in CA1 by introducing a probabilistic graphical model for the neural activity configurations in each map. Graphical models, in which a functional connectivity network accounts for the pairwise correlation structure between neuronal firing events in the recorded population (MacKay, 2003), have been applied to various areas so far (Cocco et al., 2017; Stevenson et al., 2008), e.g. to estimate the information conveyed by (Tka**c**ik et al., 2010) or the activity of (Pillow et al., 2008; Schneidman et al., 2006) retinal ganglion cells in the presence of visual stimuli, to detect learning-related changes in functional connectivity in the prefrontal cortex (Tavoni et al., 2015, 2016).

We apply our graphical-model decoder to already published (Jezek et al., 2011) and some new recordings of the hippocampal activity in CA1, performed within the teleportation setup of Jezek et al. (2011). Our decoder shows very good performances in terms of precision and statistical properties in CA1. It allows us, in particular, to identify transitions between spatial representations in CA1 in a statistically robust way. Remarkably, we find that the frequency of these flickering events is increased even minutes after a teleportation switch.

It is important to stress that, in contradistinction with previously used map decoders, ours does not use any position information. It can therefore be applied to decode and study the dynamics of general brain states with unknown input correlates, the only working hypothesis being that we dispose of reference sessions to build statistical models of the corresponding internal states.

## 2 Materials and Methods

### 2.1 Experimental methods

#### Electrode preparation and surgery

Single unit neuronal activity was recorded in three adult Long Evans male rats in hippocampal subfields CA1. Rats were implanted with a “hyperdrive” allowing for an independent positioning of 16 tetrodes organized into an ellipsoid bundle. Tetrodes were twisted from 17 um insulated platinum-iridium wire (90% and 10%, respectively, California Fine Wire Company). Impedance of electrode tips was adjusted by platinum plating to 120 – 250 kOhm (at 1 kHz). Anesthesia was introduced by placing the rat into a plexiglas chamber with seal top filled with isoflurane vapour. Then the animal was shaved and placed into the stereotaxic frame and continued the isoflurane delivery with a face mask. Breathing, heart action and reflexes were monitored continuously. Hyperdrive was then implanted above the right dorsal hippocampus at coordinates AP 3.8 mm and ML 3.2 mm relative to bregma. Stainless steel screws and dental acrylic were used to stabilize the implant on the skull. Two of the screws served as the hyperdrive ground.

#### Tetrode position

The tetrodes were slowly approached towards CA1 or to CA3 within 2-3 weeks after the surgery while the rat was resting in a comfortable pot on a pedestal. To maintain stable recordings, electrodes were not moved at all before and during the experiment on a given day. The recording reference electrode was positioned in corpus callosum. Additional reference for EEG was placed in stratum lacunosum moleculare.

#### Recording procedures

Neural activity was recorded while the rat was behaving in an apparatus described by Jezek et al. (2011). Signal was recorded differentially against the reference tetrode. Hyperdrive was connected to a multichannel, impedance matching, unity gain headstage and its output conducted through a 82-channel commutator to a Neuralynx digital 64 channel data acquisition system. Signal was band-pass filtered at 600 Hz–6 kHz. Unit waveforms above individually set thresholds (45-70 *μ*V) were time-stamped and digitized at 32 kHz. Position of the light emitting diodes on the headstage was tracked at 50 Hz to assess the animal’s position. For the purpose of this study only data from intervals when the rat’s movement speed exceeded 5 cm/sec were used. Broadband EEG from each tetrode was recorded continuously at 2000 Hz.

#### Spike sorting and cell classification

Spikes were sorted manually using 3D graphical cluster-cutting software (SpikeSort, Neuralynx) The feature space consisted of three-dimensional projections of multidimensional waveform amplitudes and energies. Autocorrelation and crosscorrelation functions were used as additional separation tools. Putative pyramidal cells were distinguished from putative interneurons by average rate, spike width and occasional complex spikes.

#### Histology

After the experiment was finished, the rat was overdosed with a barbiturate and was perfused intracardially with saline followed by 4 % formaldehyde. Brain coronal sections (30 *μm*) were stained with cresyl violet. Traces of all 14 tetrode locations were identified. Each tip location was considered as the place in the section before the tissue damage became negligible. Only recordings from tetrodes with their tips in CA1 were used in this study.

#### Behavioral procedure

Animals were first pre-trained according to the procedure described in Jezek et al. (2011). Briefly speaking, the apparatus consisted of two identical black plastic boxes (60 *×* 60 cm, 50 cm in height). The two environments differed only by sets of light cues, one placed on the upper rim of the box, the second was positioned under the semi-transparent floor with an additional cue on one wall, respectively. There were no other visual cues present as the experiment was otherwise carried in darkness provided by surrounding light-proof curtains. The training consisted of four phases. Initially, the two boxes were connected with an alley so the rat could freely explore both of them within three 20 min. sessions for 3 days. In the second phase, after the first 20 min. session, the alley was removed and the animal was placed into box *A* or *B*, respectively, in a quasi-random manner so that it received two 10 min. sessions in each of them, respectively. The next day the rat received two 10 min. sessions in each environment as the day before. Then we removed the double maze and replaced it with a single box equipped with both sets of lights that was presented at the original locations with just one cue set switched on at the given session. The rat was given another two 10 min. sessions in each environment that day. Finally, the next day, after two sessions in the original locations, the box was presented in a central location. Again, the animal was presented another two 10 min sessions in each environment, respectively, in a quasi-random order. In all stages, the running sessions were separated by a 20 min. break in the resting pot. On the test day, both environments were presented in two “reference” recording sessions (10 min each). After a 20 minutes break, the test session begun. The animal was inserted to the box with one set of lights on, and the lights were switched between the both sets after couple of minutes of recording.

### 2.2 Data structure

#### Cross validation of environment decoding methods

For the validation of environment decoding methods (Section 3.2) a total amount of four recording sessions were used. Two of them, one in the environment *A* and one in the environment *B*, called *reference* sessions, were used to infer activity models and reference statistics. The other two (again one in environment *A* and one in environment *B*) have then been used as *test* sessions, *i.e.* to assess the performance of our method for decoding which environment is internally-represented by the rodent.

#### Teleportation sessions

In the post-teleportation analysis shown in section 3.3 we used recordings from three experiments performed in three different animals (one of them was already used in the original Jezek et al. (2011) study). Each data set included two reference sessions for both environments and one or two teleportation sessions, each containing one single light switch. The switch between light cues was in total performed four times (direction balanced, *A* to *B* or vice versa), and the activity was recorded for some minutes before and after the teleportation.

### 2.3 Map decoding methods

We consider two classes of decoders: *Rate-map based decoders*, which expressly use the knowledge of place fields and the rat trajectory as an input, and *Activity-only decoders* that do not rely on any information about the correspondence between position and neural firing. Throughout this section neural activities are binned with time resolution Δ*t*; we define the number of spikes of neuron *i* in time bin *t*, *n*_*i,t*_, and the binary activity, *s*_*i,t*_ = min(*n*_*i,t,*_ 1). Little information is lost when considering *s* instead of *n* as long as Δ*t* is smaller than the typical inter-spike interval of the cells.

#### Activity-only decoders

##### Bayesian approach to map decoding

We introduce probabilistic models for the distribution of activities {*s*_*i*_}_*i* =1*…N*_ in a time bin, *P* ({*s*_*i*_}, Θ). Those models are parametrized by a set of variables, Θ, which are fitted to maximize the likelihood of the data in reference sessions. Two sets of parameters Θ(*m*) are fitted, one for each reference session *m* = *A, B*. We then define the difference in log–probabilities

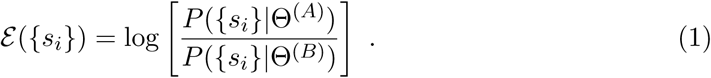

The sign of the quantity *ε*({*s*_*i,t*_}) may be used to decode the map in time bin *t*. Significance levels, based on the percentiles of the distribution of *ε* can be imposed, see Results, Section 3.3.

##### Independent-cell model

The simplest way to model the firing properties of the neural population is to assume that the neural activities *s*_*i*_ are independent from cell to cell. For each map *m*, the probability distribution *P* is parametrized by a set 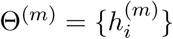 of *N* ‘inputs’ 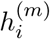:

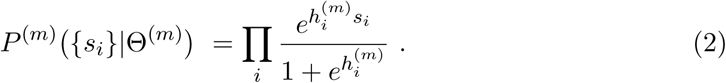

Each input parameter is fitted in order to match the average value of *si* with *P* (*m*) and the mean value 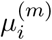 of *s*_*i,t*_ across the time bins *t* in reference session relative to map *m*. This procedure yields 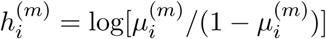.

##### Graphical Ising model

A more accurate probabilistic model for the activity of the cell population is obtained when pairwise correlations between neural activities *si* in a time bin are taken into account. For each map *m*, we introduce couplings 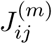 to express the conditional probability that cell *i* is active given the activity of cell *j*. The probability distribution *P* (*m*) is now parametrized by the set 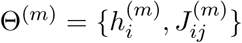 of *N* inputs 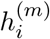 and 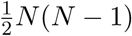 couplings 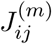:

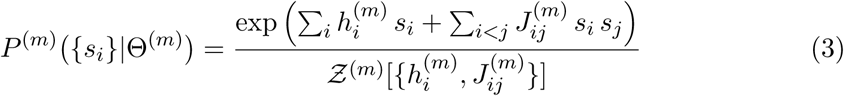

where *Ƶ*^(*m*)^ is a normalization constant. Parameters *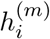* and 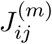 are computed to match the average values of *s*_*i*_ and *s*_*i*_s_*j*_ with *P* and, respectively, the mean values of *s*_*i,t*_ and *s*_*i,t*_*s*_*j,t*_ across the time bins *t* in reference session relative to map *m*. This hard computational problem can be approximately solved with the Adaptive Cluster Expansion (ACE) algorithm (Barton and Cocco, 2013; Barton et al., 2016; Cocco and Monasson, 2012; Cocco et al., 2017), which provides estimates of the parameters 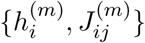 and Ƶ(*m*) in Eq. (3).

##### Adaptive Cluster Expansion (ACE)

The log-likelihood of the model parameters given the neural activities, log *P*, is regularized, *i.e.* added a term penalizing large couplings. It is expanded as a sum of contributions corresponding to clusters (subsets) of variables (Barton et al., 2016; Cocco and Monasson, 2012). Clusters of increasing sizes are recursively built from smaller clusters and added to the expansion if their contributions to the log-likelihood exceed some threshold value. The value of the threshold is iteratively decreased, until the 1- and 2-point statistics of the data are reproduced (within the expected sampling accuracy). This iterative procedure builds the simplest network (smallest number and sizes of selected clusters) able to reproduce the low order statistics of the data and avoids overfitting. Statistical error bars on the inferred inputs and coupling parameters are estimated (Barton et al., 2016). The threshold value, the number, and maximal size of selected clusters at convergence are given in Appendix A.

#### Rate-map based decoders

##### Computation of rate maps

The squared box is partitioned into a 20 *×* 20 grid of 3 *×*3 cm2 bins, and the rat position during the two reference sessions is discretized with respect to this grid. The coordinates (*x*_*t*_, *y*_*t*_) associated to time bin *t* correspond to the first spatial bin visited by the rat in the time interval [*t –*Δ*t*; *t*]. We define the average firing rate 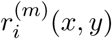 as the total number of spikes emitted by neuron *i* in the reference session *m* when the rat is at position (*x, y*), divided by the total time *T*^(*m*)^(*x, y*) spent by the animal in this spatial bin. These rate maps are then smoothed to fill missing bins through discrete cosine transform (Garcia, 2010).

##### Pearson decoder

The observed firing pattern at time *t*, *{n*_*i,t*_}_*i*=1*…N*_, is compared to the average firing rates in map *m*, 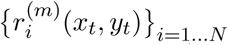, in the position (*x*_*t*_, *y*_*t*_) occupied by the animal at the same time (Jezek et al., 2011). This comparison is made through the Pearson correlation

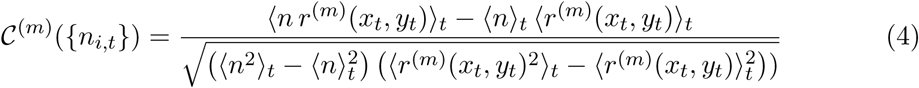

where the notation 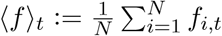 denotes the average of the quantity *f*_*i,t*_ over the *N* neurons *i* in time bin *t*. The decoding of the map in time bin *t* is done according to the sign of

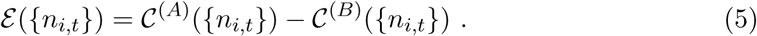

##### Dot-product decoder

The second method used in Jezek et al. (2011) compares directly the activity to the firing rates at the rat position. The decoding of the map *m* is done according to the sign of

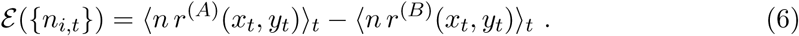

##### Bayesian Poisson rate model

This model assumes that each neuron fires independently according to a Poisson statistics, with a position-dependent firing rate 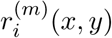 in map *m*. The probability of the number of spikes {*n*_*i*_} emitted by the neural cells in a time bin when the rat is at position (*x, y*) reads

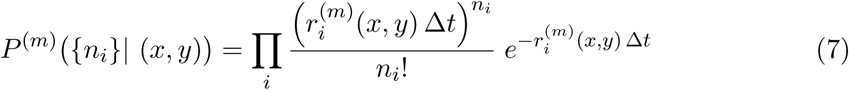

The prior probability over positions is

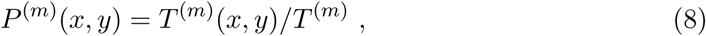

where *T*^(*m*)^ is the total recording time in reference session *m*. Assuming that both maps *m* are *a priori* equally likely, we obtain the probability of the activity conditioned to map *m* by marginalizing over positions

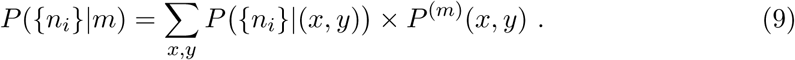

We then define the log-ratio

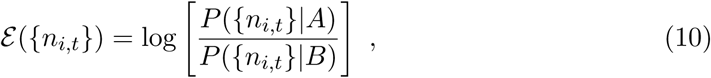

whose sign will be used to decode the map in time bin *t*.

### 2.4 Performance measure of a binary decoder

To quantitatively assess decoding performance of map-decoding methods we refer to binary classifier theory (Bradley, 1997; Hanley and McNeil, 1982; Metz, 1978; Pencina et al., 2008).

#### Receiver Operating Characteristic (ROC) diagrams

A standard framework to assess the performance of binary decoders is the so-called ROC diagram (Hanley and McNeil, 1982). For each time bin *t* the decoder outputs either map *A* or map *B*. To match the vocables used in the ROC framework we will arbitrarily say that the output is *positive* if the map is decoded to be *A*, and *negative* if the map is predicted to be *B*. If the output of the decoder matches the environment defined by the light cues at the same time *t*, the prediction is said to be *True*, otherwise it is said to be *False*. For instance, a time bin such that the decoder predicts *A*, in agreement with the cues, corresponds to a True Positive event. The 2 *×* 2 possible events are shown in Table 1. Two important quantities are: the True Positive Rate (TPR, also called Recall), that is, the number of true positive predictions divided by the total number of positive events, and the False Positive Rate (FPR), that is, the number of false positive predictions, divided by the total number of negative events. In other words, the TPR measures the fraction of time bins with *A*–cues that are correctly decoded as *A*, while the FPR is the fraction of time bins with *B*–cues that are incorrectly predicted to be *A*.

**Table 1:** Denominations used for the four possible events, depending on the output of the decoder and on the environment-defining cue. The cue is not changed throughout the reference session.

|  |  |  |  |  |
| --- | --- | --- | --- | --- |
| decoder output | $A$ | $B$ | $A$ | $B$ |
| cue | $A$ | $A$ | $B$ | $B$ |
| denomination | True Positive | False Negative | False Positive | True Negative |

Our binary decoders are all based on thresholding the *estimator* variable *ε*. Within the Bayesian framework, for instance, we compute *ε* as the difference between the logarithms of the posterior probabilities of *A* and *B*, and output Positive if the difference is larger than *θ* = 0, Negative otherwise. The value of the significance threshold *θ* can be arbitrarily changed, with the consequence of modifying the TPR and FPR values. A ROC curve shows the parametric plot of TPR vs. FPR as the threshold varies, and describes a curve in the unit square, see Results, Section 3.2. The two extreme points of the ROC curves have coordinates (0,0), and (1,1); (0,0) is obtained for a very large significance threshold *θ*, the decoder never outputs Positive and both TPR and FPR vanish; (1,1) is obtained when the significance threshold is very low, the decoder always outputs Positive and both TPR and FPR are equal to unity. Very good decoders are such that the TPR is close to unity, while maintaining a very low value for the FPR. A random-guessing decoder would give equal values for the TPR and FPR, and the ROC curve would coincide with the diagonal of the unit square.

A complementary measure of decoding performances is the Precision versus Recall (or TPR) curve, obtained by scanning the values of the significance threshold *θ*, see Results, Section 3.2. The Precision is defined as the number of true positive events, divided by the total number of positive predictions. When lowering the significance threshold the Precision decreases from 1 to 0, while the Recall increases from 0 to 1.

##### Area Under the Curve (AUC)

A quantitative measure of the decoding performances is the Area Under the (ROC) Curve (Hanley and McNeil, 1982). According to this measure, the ideal decoder has AUC = 1, while random guessing would give AUC = 0.5. Note that this measure is invariant with respect to the arbitrary choice of assigning *positive* value to environment *A*: if we assign positive to *B* and negative to *A* instead of the previous choice, ROC curves will undergo a symmetry transformation with respect to the top-left/bottom-right diagonal, resulting in an identical area under the curve. This is granted by the fact that positive and negative values are mutually exclusive and complementarily cover the whole data set: for each *θ* value the fraction of False Positive (*B* decoded as *A*) equals one minus the fraction of True Negative (*B* decoded as *B*) events.

### 2.5 Continuity prior for map decoding

A continuity prior can be included in map inference in order to reduce noise in the decoding and highlight clusters of contiguous transited time bins. To do so, we consider the output {*εt*} of the map decoder (see Section 2.3); for Bayesian decoders *t* is the difference between the log-likelihoods of the two maps *mt* = +1 and 1 in time bin *t*. We then introduce a prior, controlled by a strength parameter *K*, which favors persistence between decoded maps in nearby time bins. Informally speaking, *K* is the cost (in log-likelihood) we are willing to pay for flipping the map index in time bin *t* predicted by the sign of *t* to its opposite value, if it then matches the map indices of the neighboring time bins, *t–* 1 or *t* + 1. The prior may thus be effective in changing the map prediction *mt* if the differences between *ε*_*t - 1*_, *ε*_*t*_, *ε*_*t+1*_, … are of the order of *K* (in absolute value). Two situations are encountered: (1) for some decoders, *e.g.* Pearson, *ε*_*t*_ takes value in [ −2; 2], and the variations of *ε* over successive time bins is bounded; (2) for other decoders, *e.g.* Independent-Cell, Poisson and Ising, the difference between *ε*_*t*_ and *ε*_*t+1*_ can take arbitrarily large values and show wide fluctuations as *t* varies across the recording. In the latter case, a uniform prior *K* is unadequate in large portions of the recording. To circumvent this difficulty we introduce a scale factor *β <* 1, and multiply all outcomes *t* by this factor. As a result we get a smoother time course of *εt* over the time index *t*, on which a uniform prior can now be applied.

The joint probability of the time sequence of map predictions {*m*_*t*_} reads

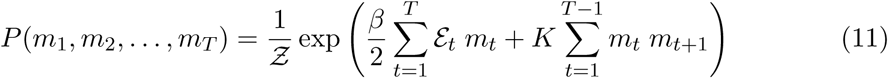

where Ƶ is a normalization coefficient. To decode the map in time bin *t* we compute the marginal probability *P*_*t*_ over *m*_*t*_ from the joint distribution *P*. Exploiting the analogy with the one-dimensional Ising model of statistical physics, this computation can be done with the transfer matrix method, also called dynamic programming, in a time scaling linearly with the total number of time bins. Then the outcome of our combined decoder+prior is

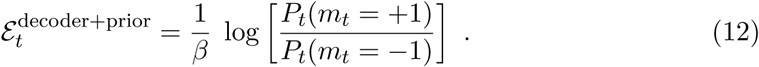

The presence of the 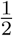 and 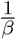 factors in, respectively, Eq. (11) and Eq. (12) ensure that, for 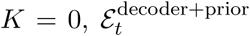 and *ε*_*t*_ coincide. In practice we choose 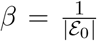, where *ε*_0_ : = max_*t*_ {|*εt*|}.

#### Induced correlation as a function of *K*

The transfer matrix technique allows us to compute also the correlation between the maps decoded *τ* bins apart, defined as

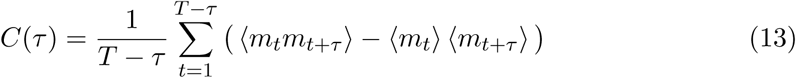

where the angular-bracket notation denotes the average over the probability distribution in Eq. (11). *C*(*τ)* decays exponentially with *τ*, over a characteristic ‘time’ monotically growing with *K* in Eq. (11), see Results, Section 3.2.

## 3 Results

### 3.1 Decoding methods and number of parameters used

We start by presenting map-decoding methods and their performances. For each environment, *A* and *B*, we have two recorded sessions with constant light cues: the first one, called reference session, is used to infer the decoder parameters. The second one, called test session, is used for cross-validation, *i.e.* to assess the performances of the decoder. We compare the performances of five different decoders, described in Methods, Section 2.3. Our decoders mainly differ by the fact that they may use or not knowledge of the rat positions and of the spatial rate maps (place fields). They are also based on simple comparison methods or on more sophisticated probabilistic frameworks.

**Rate-map based decoders** require the computation of the rate maps during the reference session. Knowledge of the position 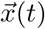 of the rats and of the neural firing rates *r*_*i*_(*t*) as a function of time *t* allows one to build the rate maps, that is, the average firing rate of each cell *i* as a function of the rat position 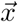, 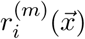 for environment *m* = *A, B*. The similarities between those reference population activities and the activity measured during the test sessions may then be used as a simple estimator of the map retrieved by the rodent. We consider two such comparison-based approaches, called *Dot Product* and *Pearson* (Jezek et al., 2011). A more sophisticated decoder, called *Poisson*, consists in assuming that each place cell *i* fires with a Poisson process, with average rate 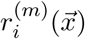 when the rodent is at position 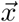, and in estimating the likelihood of the test spiking activity with this multiple Poisson process and for maps *m* = *A* and *B*. The posterior distribution for the (binary) map variable *m* can then be computed, and we decode the map as the one with larger posterior probability. Poisson is based on a more solid probabilistic framework than Dot Product and Pearson, while making use of the same rate maps estimated from the reference sessions.

**Activity-only decoders** do not need any information about rat position and place fields. Those models provide approximate expressions for the probability distribution of population neural activity over short time bins, *i.e.* of binary (silent or active neuron in the time bin) strings of length *N* (the number of recorded neurons). The *independent-cell* model is the simplest maximum-entropy model (Jaynes, 1957); it reproduces the *N* average activities of the neuron only. The second model, called *Ising* in statistical physics, is a graphical model that, in addition, reproduces the pairwise correlations between the neural activities in a time bin (Cocco and Monasson, 2011; Jaynes, 1957; Schneidman et al., 2006). The Ising model requires the inference of pairwise effective couplings between every two cells, which we have performed with the Adaptive Cluster Expansion method (Barton et al., 2016; Cocco and Monasson, 2011). Similarly to Poisson, the independent-cell and Ising models provide estimates of the likelihood of the population activity in a time bin, and can be used to compute the posterior distribution for the map variable, *m*, and to decode the retrieved map through maximization over *m*.

As a consequence the numbers of parameters to be learned from the reference sessions vary a lot with the decoders. For *N* recorded neurons (38 in one of the data sets studied here, see Materials) and a discretization of the environment into *S* (=20 *×* 20 in the present analysis) spatial bins, the numbers of parameters to be extracted from the reference sessions are, respectively *N* = 38 for the independent-cell decoder, 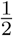 *N* (*N* + 1) = 741 for the Ising decoder, and *N × S* = 15, 200 for the Poisson, Pearson, and Dot Product decoders.

### 3.2 Cross validation of map-decoding methods

#### Inferred Ising couplings are fingerprints of environment representation in CA1

As a result of rate remapping taking place in CA1 (Fig. 1) the populations of active cells in the two environments are similar. This property can be seen from the comparison of the inputs {*h*_*i*_} in the Ising models inferred in the reference sessions of the two environments, see Fig. 2. The input *hi* to place cell *i* takes similar values across the environments; its value is indicative of the average firing rate of the cell (Methods, Section 2.3).

**Figure 2:**
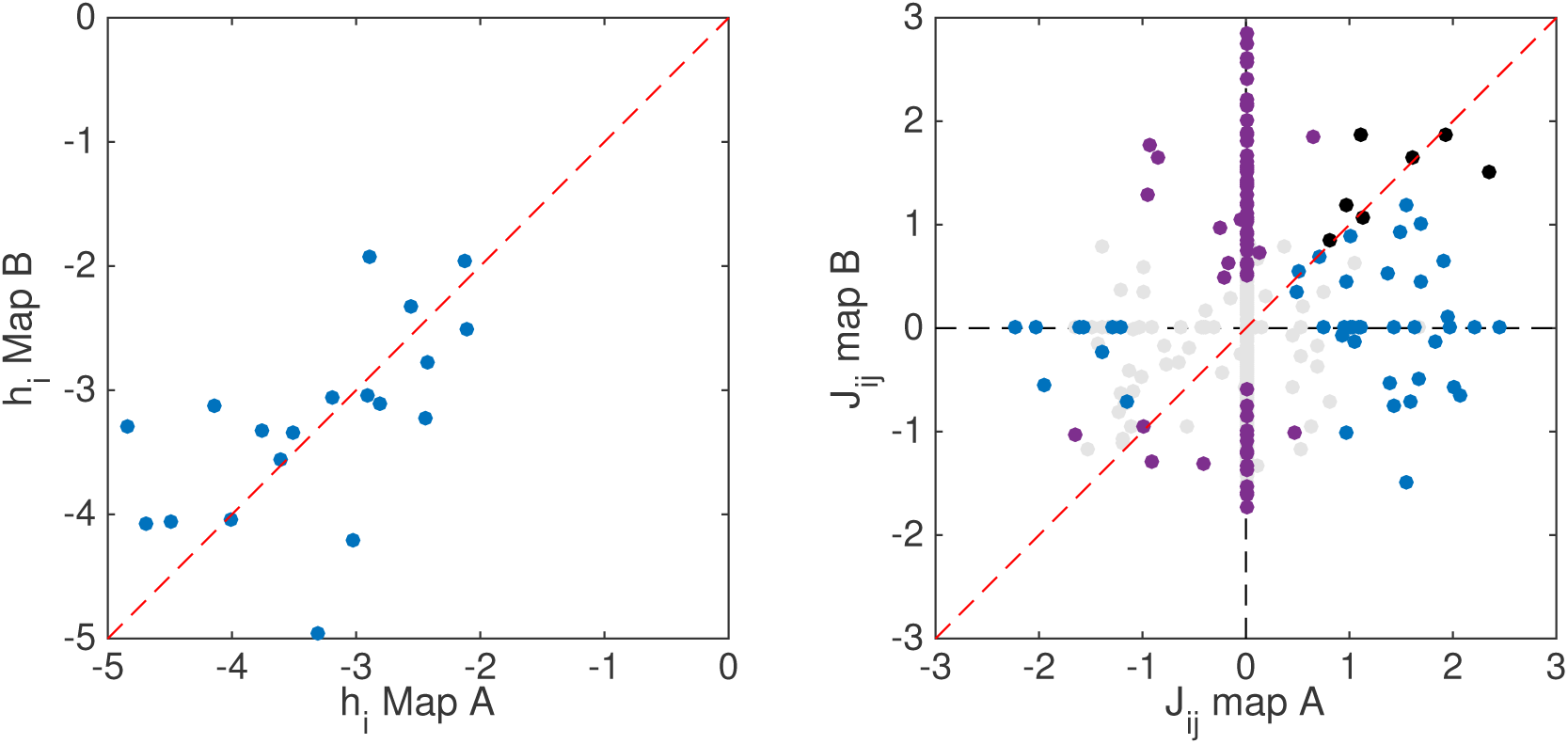
Comparison of inferred Ising parameters across the two maps. Top: Inputs *hi* of the Ising models inferred from reference sessions. Only values greater than 5, corresponding to a firing rate of c.a. 0.05 Hz in the independent-cell model, are shown. Bottom: Couplings *Jij* of the Ising models inferred from reference sessions. Dots are colored with reference to their relative statistical error (due to finite sampling) 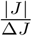: Unreliable couplings, *i.e.* such that 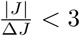 in both maps, are shown in grey (note the presence of many zero couplings produced by ACE). Couplings that are reliable only in one map are shown with purple (A) and blue (B) dots. Couplings reliable in both maps are shown in black. Analysis performed with discretization time bin Δ*t* = 120 ms.

Distinction between the neural representations of the environments in CA1 can, however, be drawn from the correlational structure of firing events in the place-cell population. Place cells with overlapping firing fields in one environment are indeed more likely to be simultaneously active during the animal’s exploration, and their activities are thus correlated. Due to remapping the amplitudes of these correlations are specific to each environment. The inferred Ising couplings {*J*_*ij*_}, which capture the direct correlation between cells *i, j* not mediated by other recorded cells (Methods, Section 2.3), are different from one environment to the other, as shown in Fig. 2.

The set of effective couplings {*J*_*ij*_} is therefore a *fingerprint* of the environment (Okatan et al., 2005), which we can exploit to distinguish between maps, *i.e.* to decode the neural representation. Note that these effective, functional couplings are not directly related to the physiological synaptic interactions, which are not accessible from the data.

Pairwise correlations are also environment specific, see Appendix B, Fig. 9. However, inferring effective couplings allows us to *score* any configuration of the population activity, that is, to quantitatively assess its similarity with typical activities in each environment, as shown below. This score is, in practice, given by the Ising probability, see Methods, Eq. (3), and heavily relies on the inferred couplings and inputs. Scoring is not possible from the knowledge of the mean activity and pairwise correlations.

#### Comparison of performances of map-decoding methods

We present a systematic study of the performances of map-decoding methods in CA1 within the framework of binary-decoder theory, see Methods, Section 2.4 for a detailed description. Results are reported in Fig. 3.

**Figure 3:**
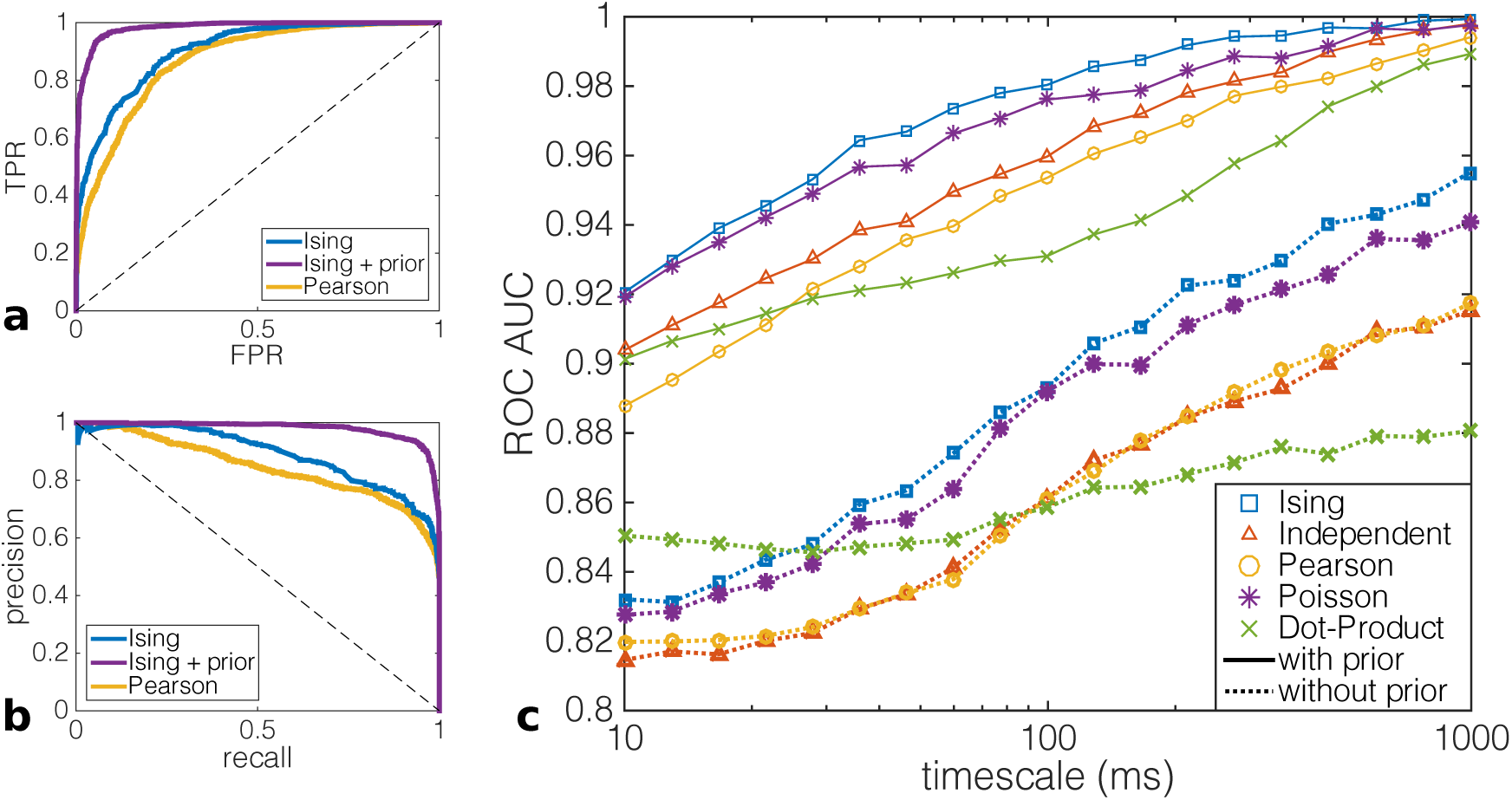
Performances of spatial representation decoders. ROC **(a)** and Precision-Recall **(b)** curves computed at fixed time scale Δ*t* = 120 ms for a combination of two test sessions in environments *A* and *B*, recorded in CA1. Maps *A* and *B* correspond, respectively, to positive and negative predictions, see Table 1. The True Positive Rate, also called Recall, is the number of true positive predictions divided by the total number of positive events. The False Positive rate is the number of false positive predictions, divided by the total number of negative events. Precision is defined as the fraction of identified positive events that are true positives. **(c)** performances of *Ising*, *Independent-cell*, *Poisson*, *Pearson*, and *Dot Product* decoders (with and without the addition of a continuity prior) as functions of the discretization time scale Δ*t*, applied to CA1 neural recordings. Full and dashed curves correspond to predictions, respectively, without and with continuity prior; in the latter case the correlation *C* in Eq. (13) decays over *t*0 = 2 time bins (Methods, Section 2.5 and Fig. 5 (a)).

We plot in Fig. 3 (a) the Receiver Operating Characteristic (ROC) curve for the Ising and Pearson decoders. Briefly speaking, ROC curve shows the value of the True Positive Rate (fractions of time bins in reference session for environment *A* for which the decoder rightly decodes map *A*) as a function of the False Positive Rate (fractions of time bins in reference session for environment *B* for which the decoder erroneously recognizes map *A*). A random decoder would have equal values for TPR and FPR, and lies on the diagonal line of the unit square in Fig. 3 (a). A perfect decoder would always recognize map *A* in environment *A* and never in environment *B*, and would thus correspond to TPR = 1, FPR = 0. Varying the threshold for significance of the decoder changes both the values of TPR and FPR, with the resulting ROC shown in Fig. 3 (a). We observe that the Ising decoder shows much better performances than the Pearson decoder. An alternative representation of the decoder performances is given by the Precision-Recall curve, shown in Fig. 3 (b), see Methods, Section 2.4 for definition.

A measure of the accuracy of the decoder is given by the integral of the ROC curve, called Area Under the Curve (AUC), which ranges from 0.5 for a random decoder to 1 for a perfect decoder. To compare the five decoders we plot in Fig. 3 (c) their AUC values as a function of the elementary time bin Δ*t*, ranging from 10 ms to 1 s.

The Ising model, which takes into account the correlational structure of the population activity, has higher decoding precision and retrieval capacity than other decoders in CA1 recordings (Fig. 3 (a,b)). As a consequence, in terms of AUC (Fig. 3 (c)), *Ising* is generally the most performant model, followed by *Poisson* and lastly by equally-performant *Pearson* and *independent-cell* decoders. *Dot Product* method is the best performant on very short time scales (< 20 ms), but its performance increases very slowly with the time bin width, and as a consequence it has the worst performance for Δ*t >* 100 ms. This behavior has an explanation in terms of sensitivity of the different models to the average number of active neurons per time bin. Bayesian models, whose predictions do not depend on the specific position of the rat at each time, rely on information conveyed by activity alone. As a consequence, when the number of simultaneously active neurons for each time bin is very small, Bayesian models may be less accurate than decoders that take into account spatial information, like Dot Product.

As a general feature we observe that the performances of all decoders improve for larger discretization time scales (Fig. 3 (c)). This result does not come from better inference of the Ising parameters, as couplings remain remarkably unchanged as Δ*t* varies, see Appendix C. The increase in performance may be simply understood as follows. Decoding performances were evaluated from the fraction of time bins in which the decoded map matched the one of the external environment evoked by the light conditions. In test sessions with stable external environment for several minutes, it is natural that merging larger portion of data results in more stable decoded maps, and, hence, in a larger fraction of correctly decoded maps. Similarly, improvement in decoding stability is obtained through the introduction of a continuity prior, which prevents switching back and forth between spatial maps in nearby time bins, see below for further discussion.

#### Performance of Ising decoder with number of recorded cells and duration of recording

We further analysed the behavior of the Ising decoder (as the most performant amongst presented methods) upon varying the number of recorded cells and the duration of the recording through subsampling the reference session data. As expected, the performance of the Ising decoder improves with the number of neurons and the duration of the reference data sets, see Fig. 4. We observe that fluctuations from subsample to subsample shrinks as the number of retained neurons increases, an effect that mirrors the heterogeneity of spatial and environment-related firing properties of single neurons. A relatively small subsample of the reference session, *e.g.* of duration *∼*1 min, suffices to compute a good estimate of the average firing rates, yielding performances similar to the independent model (Fig. 3, red curve, Δ*t* = 120 ms).

**Figure 4:**
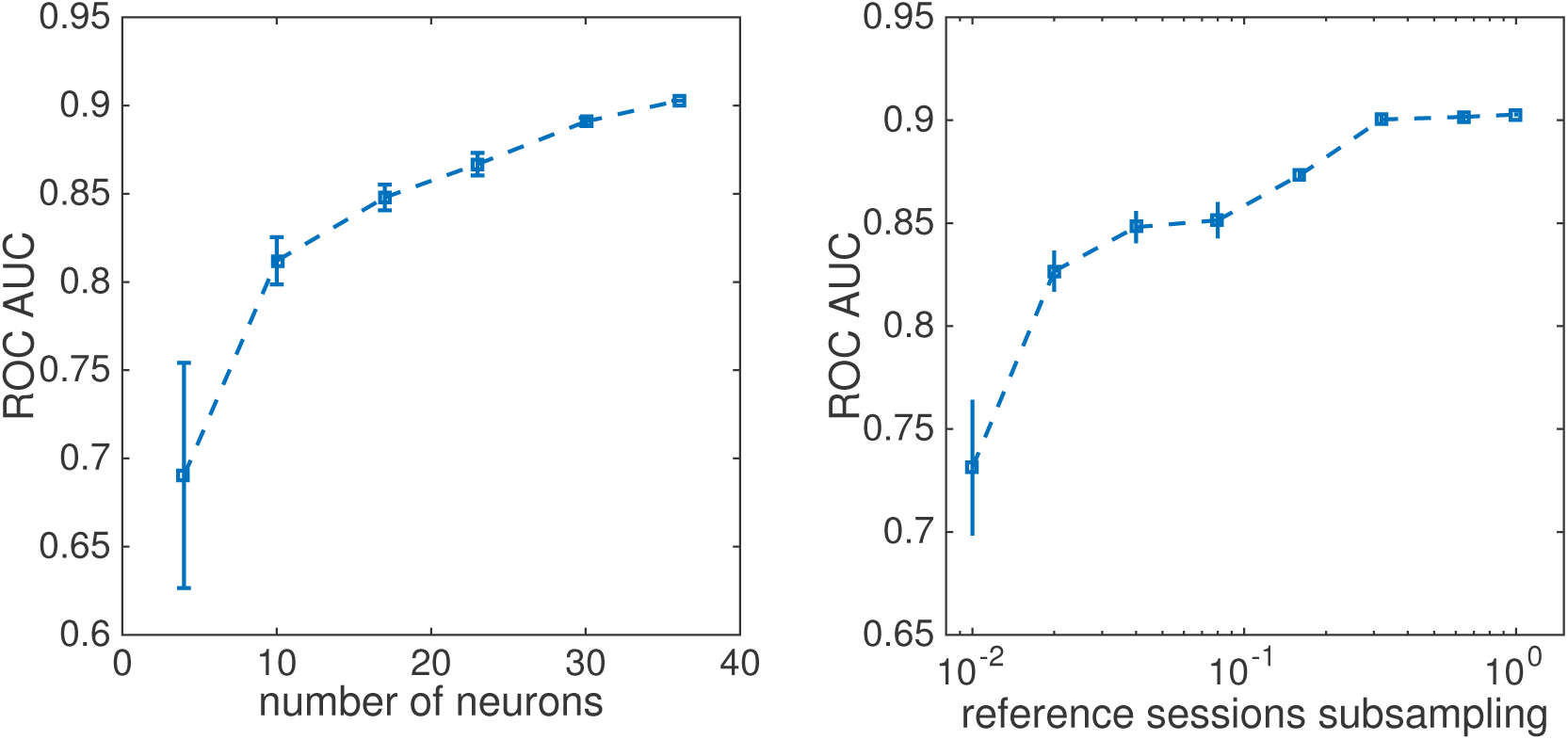
Performance of Ising decoder in subsampled conditions. Top: Performance of Ising decoder for Δ*t* = 120 ms time bins vs. number *n* of cells employed in the inference and decoding routines. For each value of *n* results are averaged over 10 randomly-chosen subsamples of cells (among the *N* = 36 recorded neurons). Bottom: Performance of the decoder as a function of the fraction of the reference session recording (subsampled from the total recordings of duration *T* = 509 and *T* = 551 seconds). For each duration considered results were averaged over 3 random subsamples of reference data.

#### Map decoding with continuity prior

Map decoding can be combined with a continuity prior that enhances persistence in the decoded maps over consecutive time bins, see Methods, Section 2.5. The motivation for the continuity prior is two-fold. First, in situations where the latency between a delivery of external stimulus and the network state change is the main parameter to be measured (e.g. after pharmacology treatment, etc.), one needs to search for a single time point of the state transition. This can be achieved by imposing a strong continuity prior, allowing for the presence of a single transition between maps along the whole recording session.

Secondly, with moderate continuity prior, dynamical events (such as state transitions) can be detected with more precision, at the price of discarding events that happen on time scales shorter than the temporal resolution set by the prior strength. To estimate this temporal resolution, we compute, for a fixed prior strength *K*, the correlation *C*(*τ)* between decoded maps in two time bins that are *τ* bins apart, see Methods, Eq. (13). This correlation decays exponentially with *τ*, see Fig. 5 (a). A persistence ‘time’ *τ*_*0*_ can be computed through an exponential fit of the correlation: *τ*_*0*_ is the characteristic number of bins over which decoded maps are persistent. Its value can be chosen at our convenience by tuning the prior strength parameter *K*, see Fig. 5 (b) and Methods, Section 2.5. Hence, we can choose a temporal resolution *τ*_*0*_ and exploit the noise-cancelling property of the continuity prior over larger time scales.

**Figure 5:**
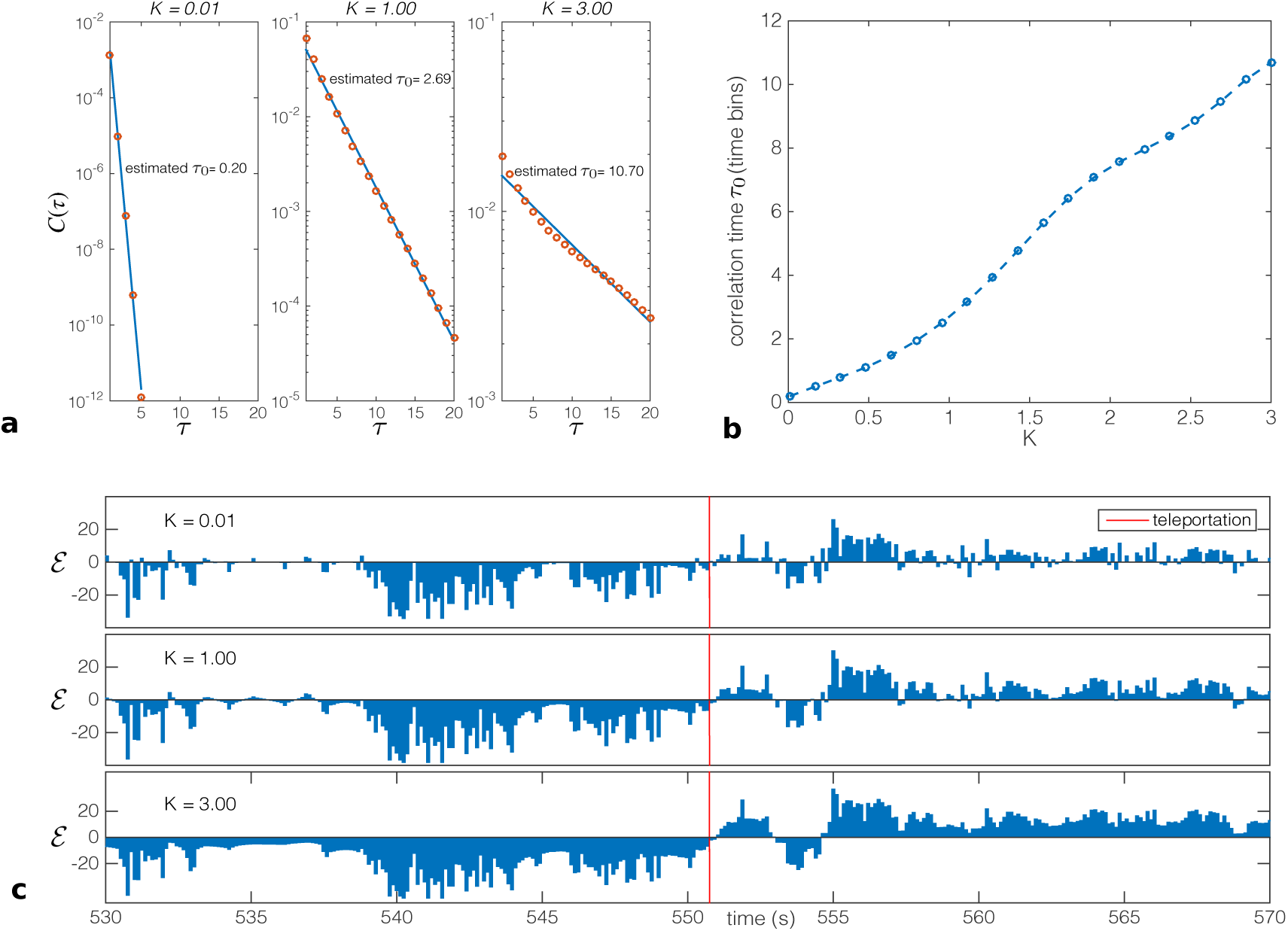
Continuity prior for map decoding. **(a)** Correlation (Methods, Eq. (13)) between maps decoded in two time bins as a function of their separation *τ* (measured in units of time bins), for three values of the prior strength *K*. Correlations are well fitted by exponential decaying functions, over a characteristic number of bins *τ*_*0*_. **(b)** Value of *τ*_*0*_ as a function of the prior strength *K*. **(c)** Application of the prior on CA1 teleportation session for different values of prior strength parameter *K*. Difference in log-probabilities of the neural activity configurations over time bons *t*. Ising decoder, with a discretization time bin Δ*t* = 120 ms.

Unless otherwise specified we set in the following the characteristic persistence time to the small value *τ*_*0*_ = 2 time bins. As shown in Fig. 3, use of this weak continuity prior enhances decoding performances with the Ising method. The AUC increases by about 10%, see Fig. 3 (c). For direct comparison, if one instead increases the time-bin resolution Δ*t* by a factor 2, the increase in AUC is much lower (Fig. 3 (c)): for instance, Ising AUC is equal to 0.90 for Δ*t* = 120 ms and to 0.92 for Δ*t* = 240 ms, while it reaches 0.98 for Δ*t* = 120 ms with a continuity prior such that *τ*_*0*_ = 2 time bins. This result shows that imposing a continuity prior is a more efficient way to reduce statistical errors in the decoding than considering larger time bins.

The observed increase in performance due to the application of the continuity prior can be explained from different perspectives. First, as explained in the Methods section and observed in Fig. 5 (a), the overall procedure introduces short-range correlations (decaying over a tunable time scale) between time bins. The resulting effect is a smoothing filter, similar to a convolution with a sliding averaging window, which acts as a noise-cancelling filter, and improves decoding precision. Secondly, the application of the prior enhances the stability of the decoded maps. This improves the decoding performance since, as pointed before, the test session is such that light conditions remain stable for long times (minutes) before the switch.

### 3.3 Transitions between maps in “teleportation” experiment

Brain hippocampal memory circuitry is a dynamic system expressing distinct states of activity - neural representations of surrounding space - with attractor properties (Colgin et al., 2010; Jezek et al., 2011; Wills et al., 2005). We applied our Ising decoder to dynamically identify those states to CA1 recordings in the ‘teleportation’ setup introduced in Jezek et al. (2011), in which the appearance of recording box is abruptly changed by switching between two familiar light cue settings (*A* and *B*, respectively) while the laboratory rat continuously explores it (Methods, Section 2.2). This procedure was shown to induce a rapid exchange of corresponding hippocampal representations in CA3, including periods of instability with spontaneous fast flickering between them. The CA1 recordings considered here include both data published in Jezek et al. (2011), and new recordings, see Methods section. Transitions between the maps were identified based on activity models of representations *A* and *B*, respectively, inferred from reference recordings in both environments under stable conditions preceding the ‘teleportation test session’.

To illustrate performance of Ising method in the post-teleportation kinetics of network state expression, we used four teleportation events recorded in hippocampal CA1 in three rats. Representative evolution of the difference in log-probabilities *ε*, see Eq. (1), of the neural activities, computed with the models inferred for the two maps from the reference sessions, is shown before and after two instances of teleportation events in Fig. 6(a). The criterion for accepting given bin as corresponding to representation of environment *A* or *B*, respectively, was set to match 1% error derived from stable reference sessions, see Fig. 6 (b). This ensures that a time bin is identified as *A* only if there is 99% (or higher) confidence that this difference in log-probabilities cannot be found in environment *B* (and vice-versa) under reference conditions.

**Figure 6:**
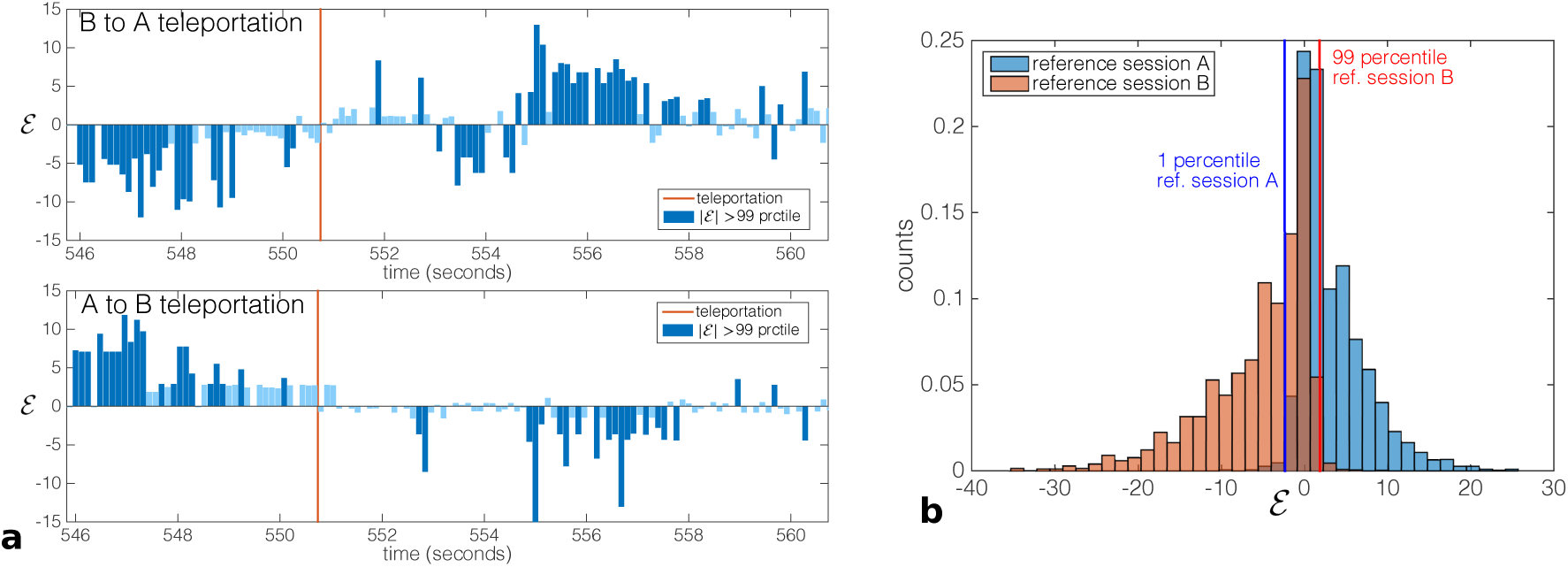
Log-probabilities of neural activities around teleportation events. Difference *ε* of the log-probabilities, Eq. (1), computed with the Ising decoder applied to the neural activity recorded in a teleportation session (**a)**, with light-cue switch from environment *B* to *A* (top) and from *A* to *B* (bottom). The light switch is marked with a red line, predictions higher than 99 percentile value of reference sessions are colored in dark blue, weaker prediction are colored in light blue. Panel **(b)** shows the distributions of differences of log-probabilities in reference sessions. A percentile value *θ* in [0, 100] (normally in the interval [90, 100]) is defined. We consider a test time bin as significantly decoded as *A* only if the log-probability difference *ε* of the activity configuration in the time bin is higher than the *θ* percentile value of reference session *B*, and as *B* only if its value is lower than the 100*- θ* percentile value of reference session *A*. The underlying reasoning is to decode a test time bin as *A* only if it is very unlikely that it comes from reference population *B*, and vice-versa.

#### Teleportation procedure induces long-term network instability

To characterize the kinetics of network state development, we identified the amount of time bins expressing a neural representation that was incongruent (non-corresponding) with the present environment, *i.e.* coding the environment presented before the teleportation. We estimated the short-term effect within interval of the first 10 seconds, and a possible long-term effect in the period that begun after 30 seconds after the teleportation has elapsed. The rates of incongruent bins are shown in Fig. 7. The amount of non-corresponding events per time bin raised from the baseline levels before the teleportation 0.013 *±* 0.002 SEM to 0.046 *±* 0.021 measured within the first 10 seconds after the teleportation (short-post effect). However, this increase was not significant, probably due to combination of large variability within the short evaluated interval (10 seconds in contrast to order of minutes of baseline state before the teleportation) and frequent empty bins (no cell active in 40.5% *±* 5.8 SEM of all bins).

**Figure 7:**
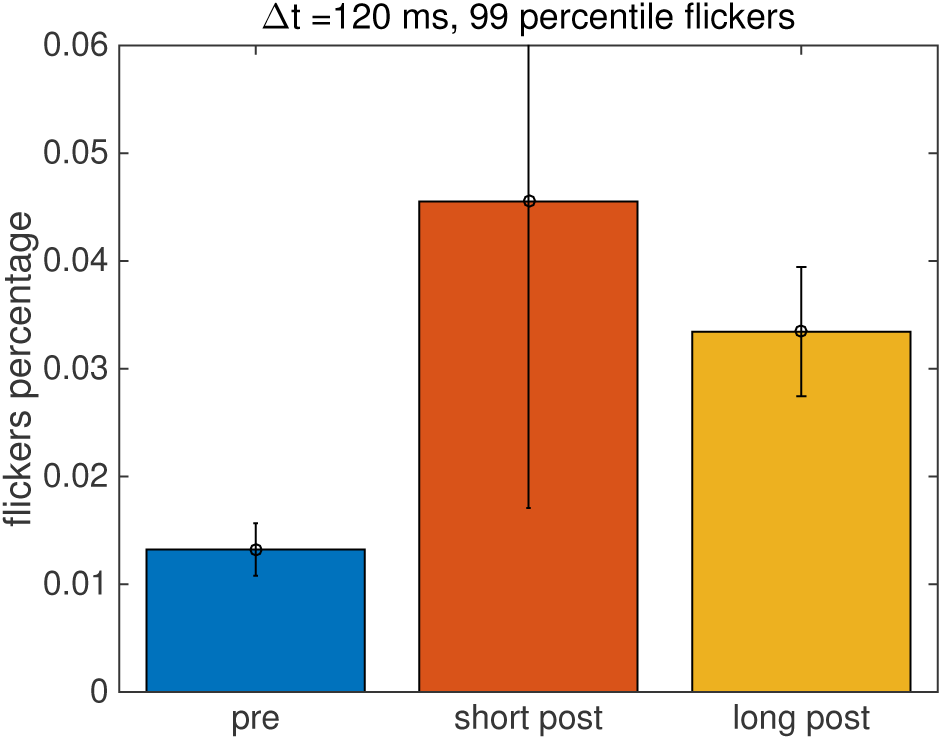
Teleportation enhances network instability over both short- and long-term periods. Percentage of temporal bins expressing the environment-incongruent coding computed in rest conditions (pre), during the first 10 seconds after a light switch (short post), and in long-term period after the teleportation event (long post, more than 30 seconds after light switch). Only bins expressing *ε*-values higher than 99 percentile of reference sessions have been taken into account; Similar results are obtained with 90 and 95 percentiles. Results were averaged over a total of four sessions recorded from three different animals. Recording durations before (pre)/after (post) teleportation equal to, respectively, 10/8, 9/9 minutes (33 cells), 12/11 minutes (17 cells), and 2.5/3 minutes (20 cells). Analysis performed with Ising environment decoder with discretization time bin Δ*t* = 120 ms.

Interestingly, the rate of flickering remained significantly increased beyond 30 seconds after the teleportation (0.034 *±* 0.021 SEM, *F* = 19.38, *p <* 0.01). Jezek et al. (2011) used temporal binning that reflected local theta oscillation (6 *-* 11 Hz) in the hippocampal circuitry. While all the results reported so far were obtained with a fixed, regular binning with a similar rate (Δ*t* = 120 ms, *i.e.* about 8 Hz), we decided to re-analyze the teleportation data in a natural theta binning as done by Jezek et al. (2011). We detected the phase of local theta oscillation based on minimum place-cell activity criterion, and the corresponding timestamps were used to define the temporal bins. We got the same pattern of results as with fixed binning (pre = 0.014 *±* 0.002 SEM, short post = 0.041 *±* 0.030 SEM, *p >* 0.05; long post = 0.030 *±* 0.005 SEM, *F* = 6.15, *p <* 0.05), yielding non-significant increase within the first 10 seconds and a significant increase after 30 seconds following the teleportation, respectively.

Last of all, we analyzed once more this teleportation data, this time with the Pearson correlation-based decoder. Neither in fixed nor theta-based binning this decoder returned significant differences between the pre teleportation and any of the post (short and long) teleportation intervals (*p >* 0.05 in all cases). This finding provides further evidence for the results depicted in Fig. 3, that is, for the better performance of Ising method over Pearson decoder for hippocampal CA1 data.

### Identification of transitions with strong continuity prior

Our network state-decoding procedure with continuity prior can be used to detect internal state-shifts under predefined criteria. For instance, when the prior strength is brought to extreme values the decoding procedure discards the fast instability-driven dynamics and, instead, returns a single state transition time point that reflects the evolution of log-likelihood values across the continuum of temporal bins. Taking as an illustration the CA1 teleportation session in Fig. 8, we see that the response of network activity state to the teleportation event is identified with high accuracy. This is a valuable tool to measure the most probable moment of network remapping even under widely fluctuating dynamics.

**Figure 8:**
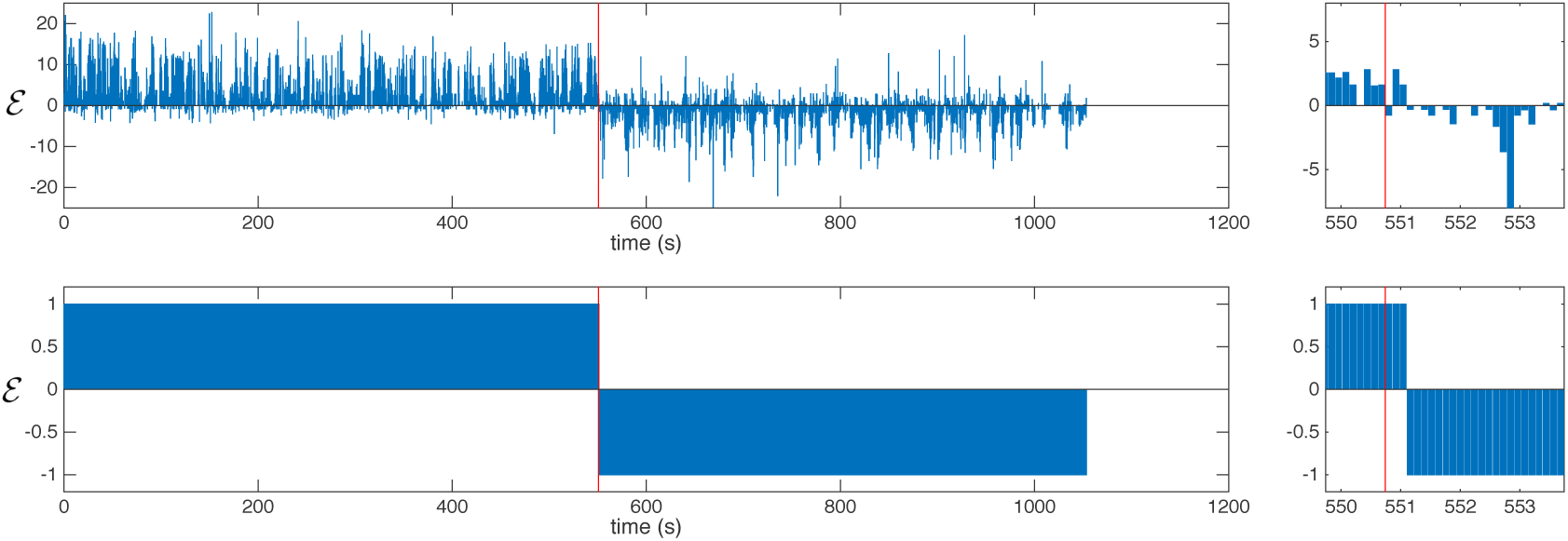
Network state transitions identified by implementation of continuity prior. Ising decoder and Viterbi algorithm with strong continuity constraint applied to neural activity in a CA1 teleportation session with enlarged examples. Light switches are marked with red lines. Analysis performed with time bin Δ*t* = 120 ms.

## 4 Discussion

### Graphical models for brain state identification in the absence of input correlates

Methods for decoding spatial representations considered in this work can be divided in two classes, depending on whether they make use of positional information or not. Remarkably, the latter methods do not show worse performances than the former approaches. In CA3, efficient decoding does not require the use of sophisticated probabilistic models: due to the quasi-orthogonality of maps, the simple independent-cell decoder, which compares the activity at any time to the average activities in environments *A* and *B* irrespectively of the rat position, shows very good performances (Posani et al., 2017). In CA1, the similarity in the firing fields across environments constrained us to consider a graphical model, the pairwise Ising model, which not only captures the average activity of the place cells but also their pairwise correlations. The higher performance of the Ising model, combined with the lower number of parameters involved in the inference process compared to firing field-based decoding methods, suggests that the correlational structure of neural firing activities conveys essential information about the internal representation of memorized environments.

A substantial advantage of this approach is that it can be effectively applied to other brain regions with much weaker correlation between the local activity and its inputs, *e.g.* the prefrontal cortex, or without any known input-output relation. The use of graphical models does not require any knowledge about the network inputs, as activity states are identified based on a (high-dimensional) fit of the correlation structure of the spiking data Cocco et al. (2017). The core idea, first put forward in the context of retinal data modeling (Schneidman et al., 2006), is that the model obtained after inference of the functional network is an approximate (albeit quantitatively accurate, compared to principal-component based approaches (Lin and Gervasoni, 2008)) description of the distribution of activities characterizing specifically one brain state. Provided that we have at our disposal different data sets for well-identified states (here, the reference sessions) we may later use the inferred model to decode the activity at any time. This approach has recently been applied to identify transient activation of memory-related cell assemblies in the rat prefrontal cortex (Tavoni et al., 2015, 2016). We expect, owe to its generality and its applicability to very fast time scales (down to *∼* 10 msec), further applications in future. Note that our decoding approach, based on the inference of effective pairwise couplings, could be extended to higher-order interactions. In the hypothetical situation of distinguishing between brain states that differ in high-order statistics, *e.g.* in the frequencies of 3-cell firing events only, the inference of these high-order effective interactions would be necessary to obtain an efficient decoder.

Let us also remark that, once the Ising parameters have been inferred from reference sessions corresponding to the possible states (here, maps), the computation of the log probability difference *ε*({*s*_*i,t*_}) is very fast, as it requires *O*(*N* ^2^) operations only. Our decoder could therefore be applied online, provided the neural activity configuration are available, *e.g.* through automatic spike sorting, at any time. One potential issue here is that fast spike sorting may introduce error in the activity variables, leading to wider and more overlapping distributions of *ε*, see Fig. 6 (b). Maintaining high precision in the decoding would still be possible if the confidence threshold *θ* is increased, but at a price of smaller recall, see Fig. 3 (b).

### Functional connectivity-based models for map decoding: Ising and other models

The Ising decoder introduced in this article yields the highest performance on all time scales Δ*t* in CA1 (Fig. 3). While we have here mostly considered the activity vectors as discretized in regular-spaced time windows of duration Δ*t*, our approach was also easily extended to process activity in elementary windows in correspondence to Theta cycles. It would be interesting to pursue the latter analysis to deepen our understanding of the role of Theta oscillations for the dynamics of transitions (Jezek et al., 2011) in CA1, and to assess the plausibility of the different transition scenarios (temporary disappearance or coexistence of both map representations) put forward by theoretical studies (Monasson and Rosay, 2015).

In this regard, repeating the present study with probabilistic models capable of capturing some aspects of the activation dynamics in recorded spiking sequences, such as Generalized-Linear Models (Truccolo et al., 2005), could be potentially interesting. Contrary to their Ising model counterparts effective couplings in the GLM approach are not necessarily symmetric, and may reflect specific ordering in neuron activations. However, some basic assumptions underlying GLM, such as the Poissonian nature of firing events are questionable for hippocampal place cell activity (Fenton and Muller, 1998). Another potentially interesting alternative is provided by reverse engineering of networks of Integrate-and-Fire neurons (Koyama and Paninski, 2010; Makarov et al., 2005; Monasson and Cocco, 2011), which were already applied to recordings, *e.g.* of retinal data with tens of neurons.

### Instabilities in hippocampal space representations

In the CA3 area of hippocampus, patterns of place-cell activity across different environments behave as uncorrelated network states with attractor properties (Wills et al., 2005). Transitions between those hippocampal activity states were recently studied based on recordings taken during a free exploration in two environments in an experimental paradigm shown to induce rapid switches (Jezek et al., 2011). In the present paper we used multiunit recordings from hippocampal area CA1. Both CA3 and CA1 are parts of the entorhino-hippocampal loop, an essential circuit for spatial memory and navigation in mammalian brain. Despite being directly connected in series (CA3 signalling into CA1), they very much differ in their architecture - while CA3 is organized as a recurrent network with attractor properties, CA1 has a feed forward structure - and in their connections with other brain areas involved into space representation (Knierim, 2006).

Use of the Ising model allowed us to robustly decode the memory state expressed in the CA1 network, with temporal resolution high enough to reflect natural time patterning of activity provided by local theta oscillation (ca. 6-11 Hz). We could track the network state kinetics following the sensory input switch. In agreement with previous report in CA3 (Jezek et al., 2011), we detected a high degree of flickering in CA1 following the switch.

Moreover, when analyzing the development of post-teleportation population vector activity on a long-term scale (20-60 sec), we found sustained network instability in CA1 spanning far beyond the 10 seconds interval reported in (Jezek et al., 2011), see Fig. 7. This effect is statistically significant with the Ising decoder but could not have been discovered with simpler, correlation-based methods. The presence of long-term instabtility in CA1 is rather surprising as the network usually reaches a relative stability within a couple of seconds after the cue switch (Jezek et al., 2011). An occasional delayed spontaneous flickering was described in CA3 as a result of repetitive teleportation within a short time period (every 40-60 seconds)(Jezek et al., 2011). This suggests prolonged (though rare) flickering effect might be present in both CA3 and CA1. In our data the persistent instability in CA1 came after one or two teleportation events on a given day, respectively.

What mechanism can account for this observation? The current view considers the short term (up-to 10 sec.) instability as a product of teleportation-induced conflict between a sudden change in the allothetic visual input (another environment presence) and a non-corresponding idiothetic signaling (no self-motion tracked traversal). Within couple of seconds the idiothetic input seems to reset as the rate of flickering dramatically decreases to levels close to the baseline steady state. The fact an occasional flickering is present longer both in CA3 and CA1 can have more reasons. The autoassociative character of CA3 is capable to store and express stable patterns of activity, but also to associate between different simultaneously active ensembles in the network. After teleportation, despite an attractor separation on a theta frame binning has been proved, an occasional overlap between both representations is present as well. Such brief coactivation of concurrent maps can eventually lead to their binding by collateral synapses or by detecting and learning their conjunction by CA1 (Treves and Rolls, 1994). Such a linkage could, under appropriately ambiguous or noisy input (e.g. encountering an odor mark dropped in the concurrent lighting conditions), eventually lead to a rare completion of the concurrent activity state. Other, rather speculative, possibility is that the observed activation of the other representation in CA3 and CA1 could be related to a reflection of past configuration of the external world, eventually to an expectation of another coming change of environment identity, so far of unknown mechanisms. Whatever input triggers the long-term flickering, these transient episodes do not occur during sharp wave/ripple complexes as they were present during strong theta network oscillation without any apparent increase of population activity. A further insight that is beyond the scope of this report is necessary to provide a better understanding of the origin and characteristics of long-term dynamics of transition between distinct hippocampal network states.

## acknowledgements

We are indebted to S. Rosay, who contributed to the early stage of the data analysis. We are grateful to J. Tubiana and A. Treves for useful discussions and suggestions. This study benefited from partial fundings from the CNRS-InphyNiTi INFERNEUR project and from GACR 15-20008S, Q-39 and NPU I LO1503 of the Czech Republic.

## Appendix A: ACE inference convergence details

The ACE inference procedure of Ising model parameters was applied with *L*2-norm regularization of strength *γ* = 5*/B*, where *B* is the total number of time bins (Barton et al., 2016). Details on the convergence are given in Table 2. The full code for Adaptive Cluster Expansion can be downloaded from the GitHub repo https://github.com/johnbarton/ACE/.

**Table 2:** Studied sessions (cv. = cross-validation, t. = teleportation, followed by number of teleportation and environment) number of recorded cells (*N)* and ACE parameters at convergence: threshold *θ* for cluster selection, cross-entropy (in natural log.), maximal size *KC* and number *NC* of selected clusters. The algorithms stops when the relative errors on single-neuron frequencies and pairwise connected correlations become smaller than unity (Barton et al., 2016).

| session | $N$ | $\theta(\times 10^{-3})$ | $S$ | $K_C$ | $N_C$ |
| --- | --- | --- | --- | --- | --- |
| cv. A | 36 | 0.89 | 5.31 | 7 | 390 |
| cv. B | 36 | 0.08 | 6.21 | 8 | 4789 |
| t. I-II A | 33 | 0.21 | 4.82 | 4 | 509 |
| t. I-II B | 33 | 1.4 | 4.33 | 5 | 198 |
| t. III A | 17 | 1.1 | 1.60 | 3 | 27 |
| t. III B | 17 | 0.81 | 1.76 | 3 | 34 |
| t. IV A | 20 | 0.89 | 5.45 | 4 | 117 |
| t. IV B | 20 | 0.99 | 4.51 | 6 | 1347 |

## Appendix B: Comparison of neuron activities across spatial maps

Similarly to Fig. 2 where we compare the Ising parameters inferred from the population activity in the two environments *A, B*, we show in Fig. 9 the probabilities of firing of all cells *i* (in fixed time bins with Δ*t* = 120 ms) and the pariwise correlations (defined as the probability that cells *i, j* fire together in a bin minus the product of their individual firing probabiltiies). We see that no substantial correlation is found in the pairwise statistics of cells across the two environments.

**Figure 9:**
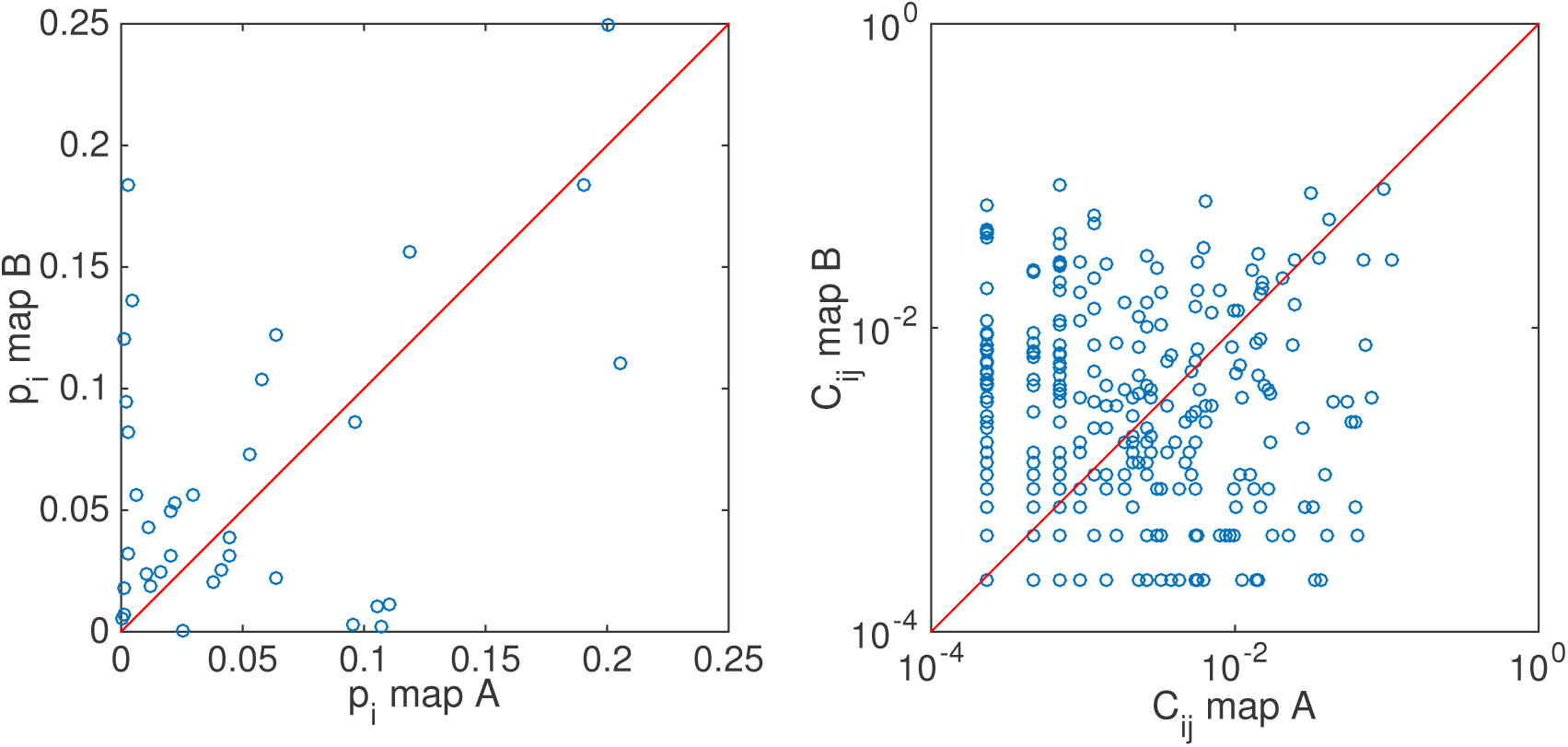
Comparison between correlations and averages of the two maps of the cross-validation reference sessions. Fixed binning with Δ*t* = 120 ms.

## Appendix C: Dependence of *J*_*ij*_ on temporal binning

Couplings inferred for time-bin duration Δ*t* = 120 ms are compared to the ones inferred for Δ*t* = 10 ms in Fig. 10. Many couplings are very similar across the two binning choices. Differences, in particular null couplings in just one of the two cases, mostly arise from sampling differences. For 10 ms time windows, it is rare to find two neurons active within the same time bin, while, for larger time bins, there is a smaller number *B* of time bins, which forces us to consider larger ACE threshold *θ*. Couplings inferred using the theta-binning discretization procedure for data are very similar to the ones inferred using a fixed time binning of 120 ms (average duration of theta cycles), see Fig. 11. A discussion of the independence of Ising couplings from the bin duration Δ*t* was done by Cocco et al. (2009).

**Figure 10:**
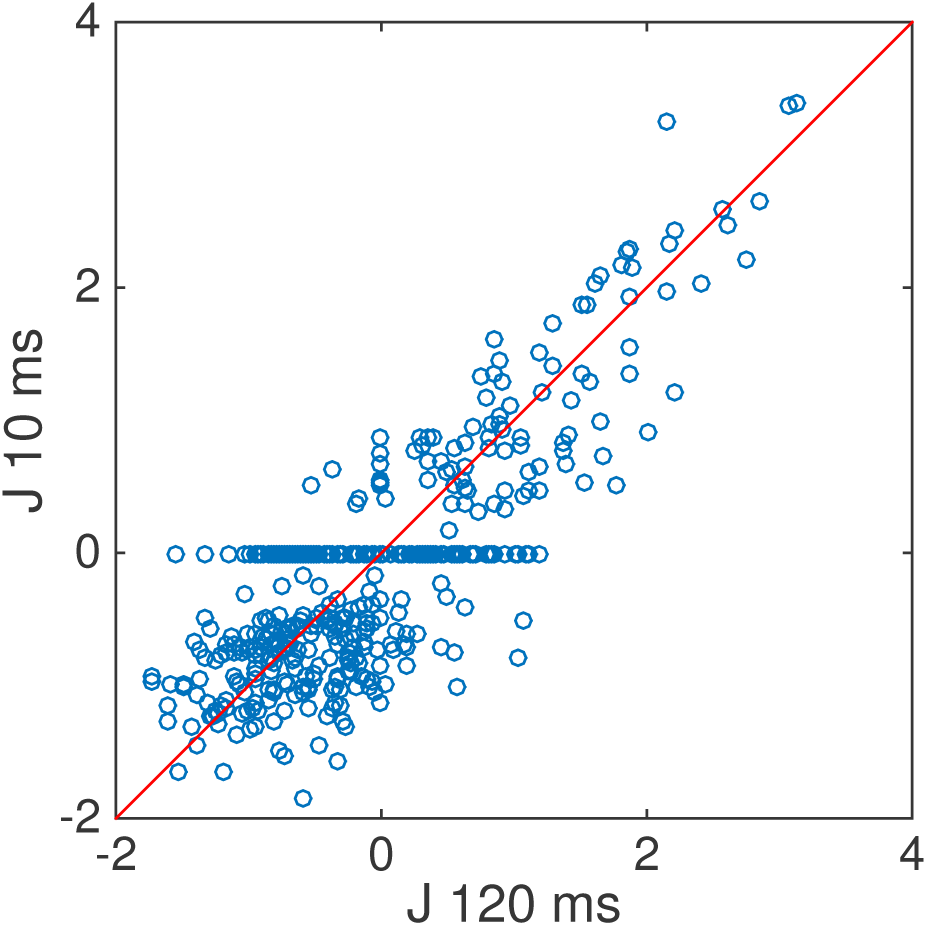
Scatter plot of couplings inferred with time bin Δ*t* = 120 ms vs. Δ*t* = 10 ms (fixed time bin discretization procedure, from cross-reference data set).

**Figure 11:**
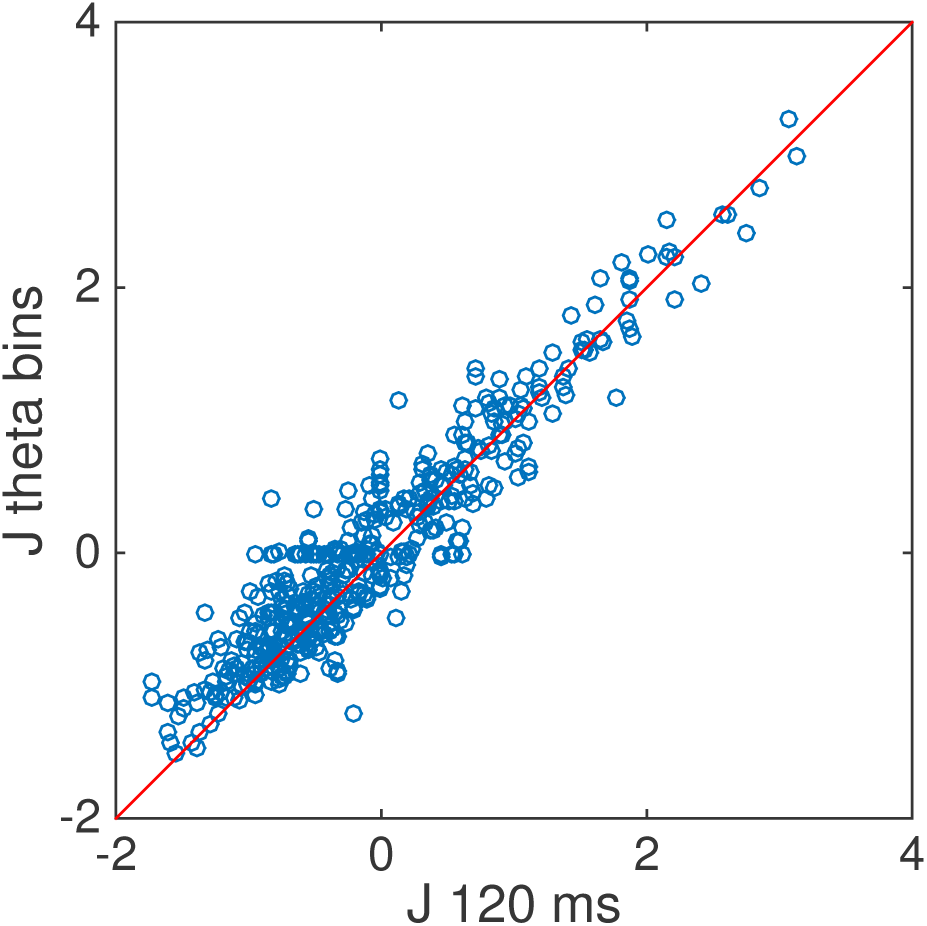
Scatter plot of couplings inferred with fixed time bin vs. theta-binning procedure (Δ*t* = 120 ms, from cross-reference data set).

